# Acquisition of cancer stem cell properties during EMT requires cell division

**DOI:** 10.1101/2021.07.01.449976

**Authors:** Petra den Hollander, Suhas Vasaikar, Maria Castaneda, Robiya Joseph, Abhijeet P. Deshmukh, Tieling Zhou, Mika Pietila, Chunxiao Fu, William F. Symmans, Rama Soundararajan, Sendurai A. Mani

**Affiliations:** Department of Translational Pathology, University of Texas MD Anderson Cancer Center, Houston TX 77030

## Abstract

Cancer cells acquire stem cell and mesenchymal properties during epithelial-to-mesenchymal transition (EMT), facilitating metastasis and chemoresistance [1-6]. In this study, we find that mammary epithelial cells quickly develop mesenchymal phenotype in response to EMT-inducing signals; however, acquiring stemness takes several days and always requires a preceding mesenchymal program. In addition, we observe that carcinoma cells, over a period of time, switch their cell division from symmetrical differentiated type to symmetrical self-renewal type. Importantly, epithelial cells can gain mesenchymal properties without undergoing cell division, but cell disivion is vital for these cells to gain stem cell properties during EMT. The EMT-induced stemness signature (SC-sig) is capable of predicting progression-free and overall-survival of breast cancer patients but not the EMT-induced mesenchymal signature (M-sig). Collectively, our findings demonstrate that the use of mesenchymal markers alone is insufficient to identify tumors with metastatic and chemoresistance potential and emphasize that the markers of EMT-induced stem cell program are central for clinical prediction. Most importantly, our data, for the first time, demonstrate that acquisition of stem cell properties during EMT depends on cell division but not the mesenchymal program.

## Chronic exposure to TGF-β1 induces a switch from symmetrical differentiated to symmetrical self-renewal type of cell division

Activation of epithelial-mesenchymal transition (EMT) using TGF-β1 has been shown to induce both mesenchymal and stem-cell properties while activating EMT [7-10]. We tested chronic exposure of TGF-β1 on MCF10A mammary epithelial cells and found that TGF-β1 gradually induced mesenchymal morphology (Fig. 1a) and increased expression of proteins associated with the mesenchymal phenotype within 2 days, such as Fibronectin, N-Cadherin and Vimentin and decreased the expression of proteins associated with epithelial phenotype such as E-Cadherin (Fig. 1b). In addition, these cells also significantly gained stem cell properties after 4 days, as observed by enhanced sphere-forming capability and aldehyde dehydrogenase (ALDH) activity (Fig. 1c, and Supplementary Fig. 1a). Comparable findings were obtained upon inducing EMT in transformed human mammary epithelial cells stably expressing tamoxifen-inducible EMT-inducing transcription factor Snail (HMLER-Snail-ER) (Supplementary Fig. 1c-e)^1^.

**Figure 1.**
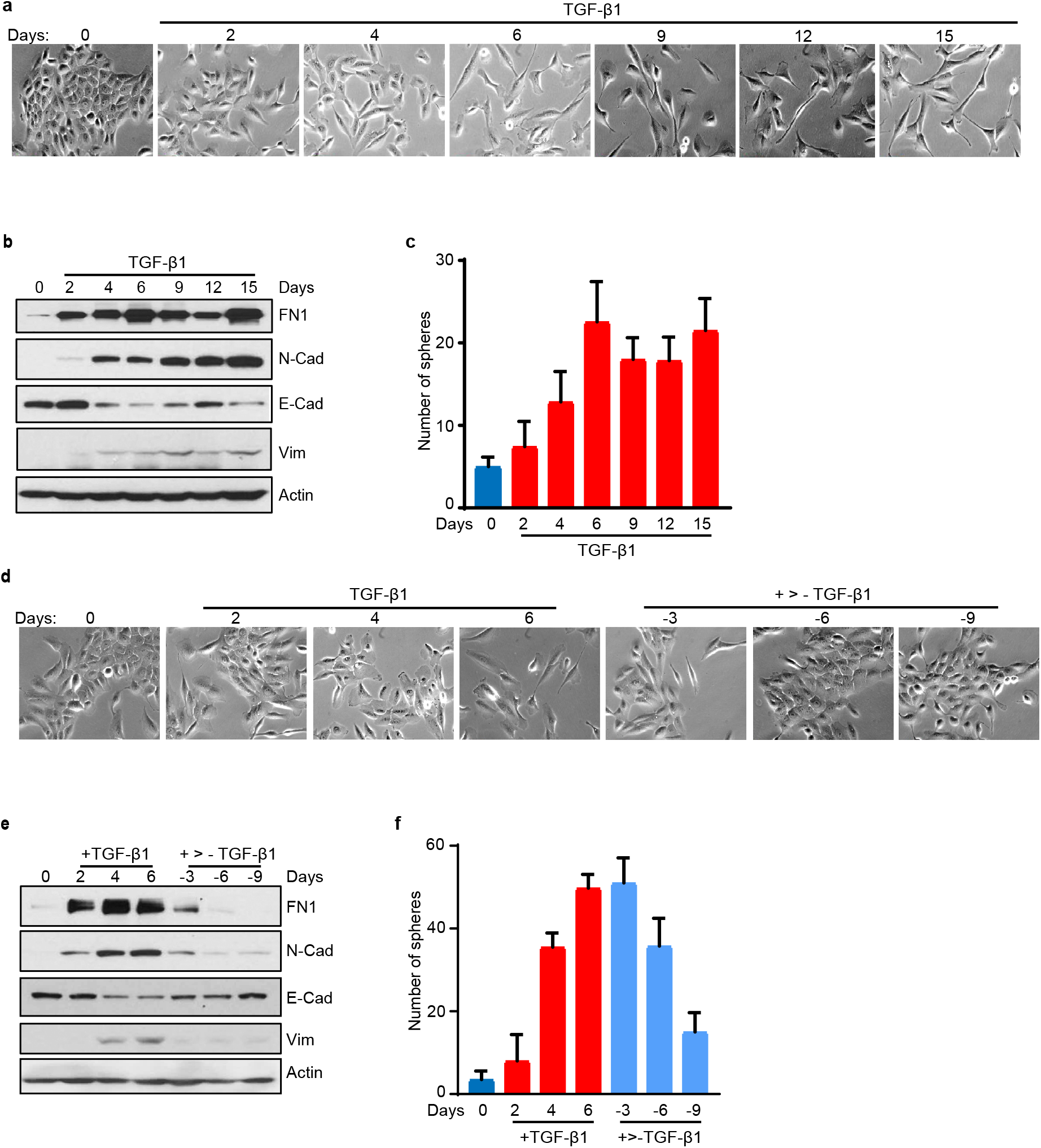
During TGF-β1 induced EMT mesenchymal properties arise before stem cell properties. **a**, Phase-contrast imaging of MCF10A cells at the indicated days during TGF-β1 exposure. **b**, Western blot analysis of indicated markers associated with EMT over 15 days of TGF-β1 exposure. **c**, Number of spheres formed by 500 MCF10A cells collected on indicated days during TGF-β1 exposure (n=4). **d**, Phase-contrast imaging of MCF10A cells during 6 days of TGF-β1 exposure and 9 days of withdrawal. **e**, Western blot analysis of markers associated with EMT on indicated days of TGF-β1 exposure and withdrawal. **f**, Number of spheres formed by 500 MCF10A cells collected on indicated days during TGF-β1 exposure and withdrawal (n=4).

In both MCF10A and HMLER-Snail-ER cells, the expression of mesenchymal markers appear within 2 days and continue to increase over a period of time (Fig 1a, b, and Supplementary Fig. 1 b and c). However, the stem-cell properties appeared only after 4 days of continuos treatment with TGF-β1 (Fig. 1c, and Supplementary Fig. 1a). Similarly, when TGF-β1 was withdrawn from the MCF10A cells, the mesenchymal morphology (Fig. 1d), and the expression of the mesenchymal markers decreased 3 days after withdrawal (Fig. 1e), while it took at least 6 days to observed a decrease in the stem cell-related ALDH activity and sphere-forming ability (Fig. 1f and Supplementary Fig. 1b, h). Likewise, when tamoxifen was removed from the HMLER-Snail-ER cells and Snail activity is no longer induced, the mesenchymal characteristics were reduced well before the stem-cell properties (Supplementary Fig. 1f-h).

Mesenchymal properties begin to appear even before the MCF10A cells undergo cell division, while 4 population doublings are required for stem-cell properties to appear after exposing to TGF-β1 (Fig. 2c). This suggests that cell division may not be required to endow mesenchymal properties but is needed to elicit stem cell properties, and most likely are acquired after multiple cell divisions. It is important to note that TGF-β1 exposure did not affect MCF10A cell proliferation (Fig. 2a), cell death (Supplementary Fig. 2a) or cell cycle phase distribution (Supplementary Fig. 2b). In addition, one cell doubling in the presence of TGF-β1 was only increased by about 3 hours relative to vehicle-treated cells (22.9 vs 19.9 hours on average respectively) (Fig. 2b) suggesting that the changes in cell death, cell proliferation, or doubling times are not the reason for the four day requirement to acquire stem cell properties.

**Figure 2.**
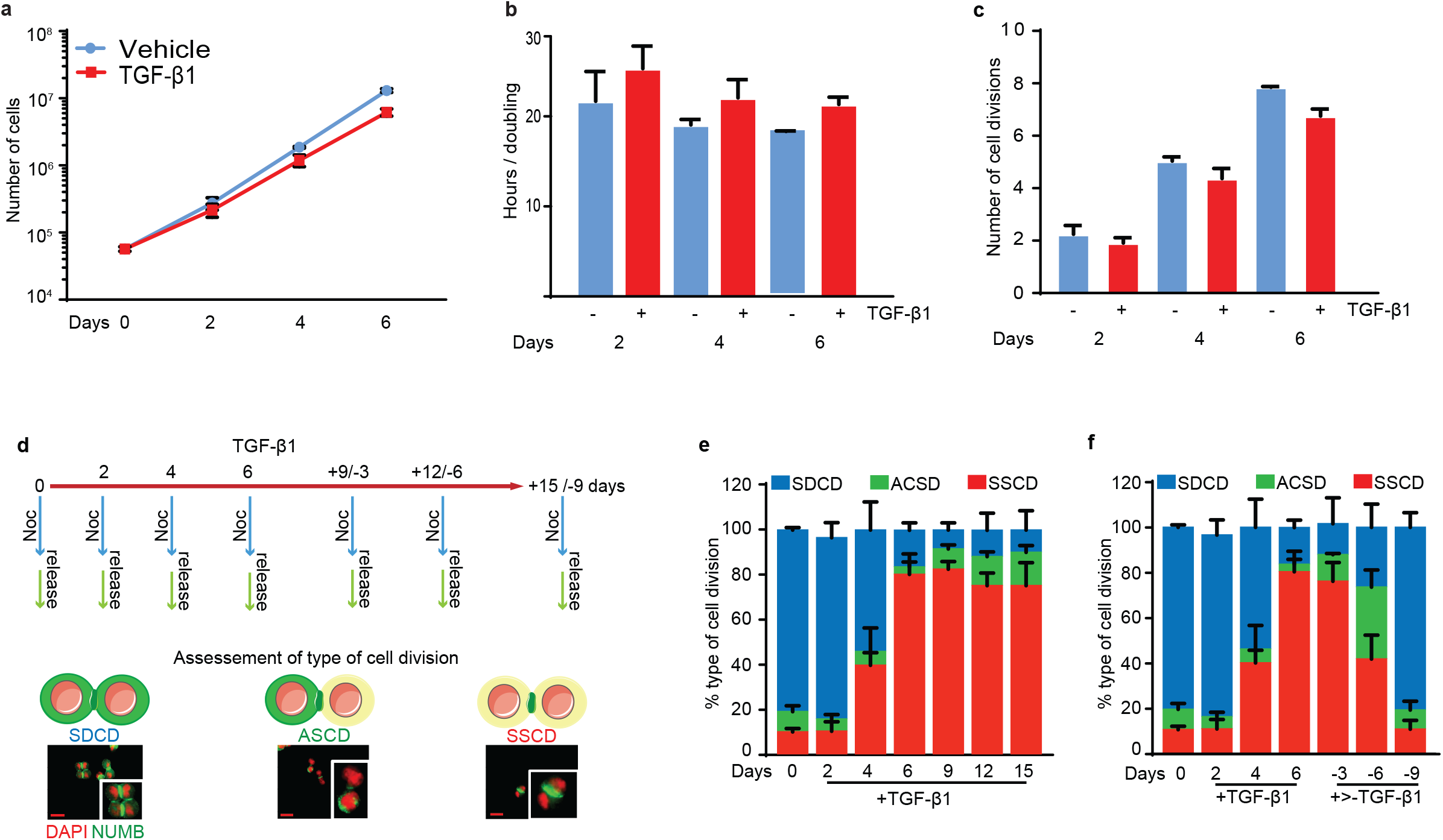
TGF-β1 induces stem cell properties by switching cell division from SDCD to SSCD. **a**, Proliferation of MCF10A cells exposed to TGF-β1 or vehicle for 2, 4, and 6 days (n=3). **b**, Population doubling time of MCF10A cells treated with TGF-β1 or vehicle for 2, 4, and 6 days(n=3). **c**. Number of MCF10A cell doublings after 2, 4, and 6 days of TGF-β1 stimulation (n=3). **d**, Schematic of immunofluorescence localization of NUMB with examples during asymmetrical stem cell division (ASCD), symmetrical stem cell division (SSCD), and symmetrical differentiated cell division (SDCD). **e**, Quantitation of cell division type during 15 days of continuous TGF-β1 exposure (n=3). **f**, Quantitation of cell division type during 6 days of TGF-β1 exposure and 9 days of withdrawal (n=3). Error bars show standard deviation.

Stem cells are known to undergo both asymmetric stem cell division (ASCD) as well as symmetric self-renewal cell division (SSCD) [11, 12], and we and others have demonstrated previously that epithelial cells acquire stem cell properties during EMT [1, 2]. Therefore, we examined whether epithelial cells during EMT switch from the regular symmetrically differentiated cell division (SDCD) to either ASCD or SSCD and the time it takes to switch the type of cell division (Fig. 2d). Surprisingly, we found that MCF10A cells, which predominantly divide via SDCD took nearly four days to change to SSCD with a maximum level of SSCD observed on day 6 (Fig. 2e and f), and remained high up to 15 days when continuously exposed to TGF-β1 (Fig.2f). HMLER-Snail-ER cells also switched their type of cell division similarly after treatment with tamoxifen to induce EMT (Supplementary Fig. 2e). Removal of TGF-β1 from the MCF10A cells or tamoxifen from the HMLER-Snail-ER cells resulted in reversal of SSCD to SDCD after 6 days of withdrawal (Fig. f). These data show that epithelial cells first acquire mesenchymal properties during EMT and after four days or five-cell divisions acquire stem-cell properties.

## Cell division is necessary to confer stem cell properties during EMT

To test the hypothesis that cell division is required to gain stem cell properties during EMT, we exposed the epithelial cells to TGF-β1 to induce EMT while simultaneously blocking cell division using thymidine (Fig. 3a). Surprisingly, regardless of the presence of thymidine and in absence of cell division, TGF-β1-exposed MCF10A cells continue to induce the expression of mesenchymal markers similar to control cells (Fig. 3b). In contrast, the cells exposed to the thymidine block did not form any stem cell-related spheres (Fig. 3d) and had significantly fewer ALDH-positive cells as compared to TGF-β1 only treated cells (Supplementary Fig. 3a). Sphere formation involves proliferation and differentiation of stem-like cells (Ref). It is conceivable that the thymidine-treated cells failed to form spheres due to the lack of cell proliferation. Nevertheless, after thymidine withdrawal, the TGF-β1 and thymidine-treated cells proliferated at similar rates as compared to vehicle treated cells (Fig. 3c). To investigate if the inhibition of stem cell properties was primarily due to a block in cell division and not due to the presence of thymidine, we treated MCF10A cells with lovastatin, which arrests cells in the G1 phase of the cell cycle. We found that blocking cell division, regardless of the inhibiting agent, prevents the appearance of TGF-β1-induced stem-cell, but not mesenchymal properties (Supplementary Fig. 3b-d). Most importantly, there was no switch of the SDCD to SSCD type of cell division in cells treated with both TGF-β1 and thymidine (Fig. 3e). Substantiating these observations, the tamoxifen-inducible HMLER-Snail-ER cell model also yielded similar results (Supplementary Fig. 3e-h).

**Figure 3.**
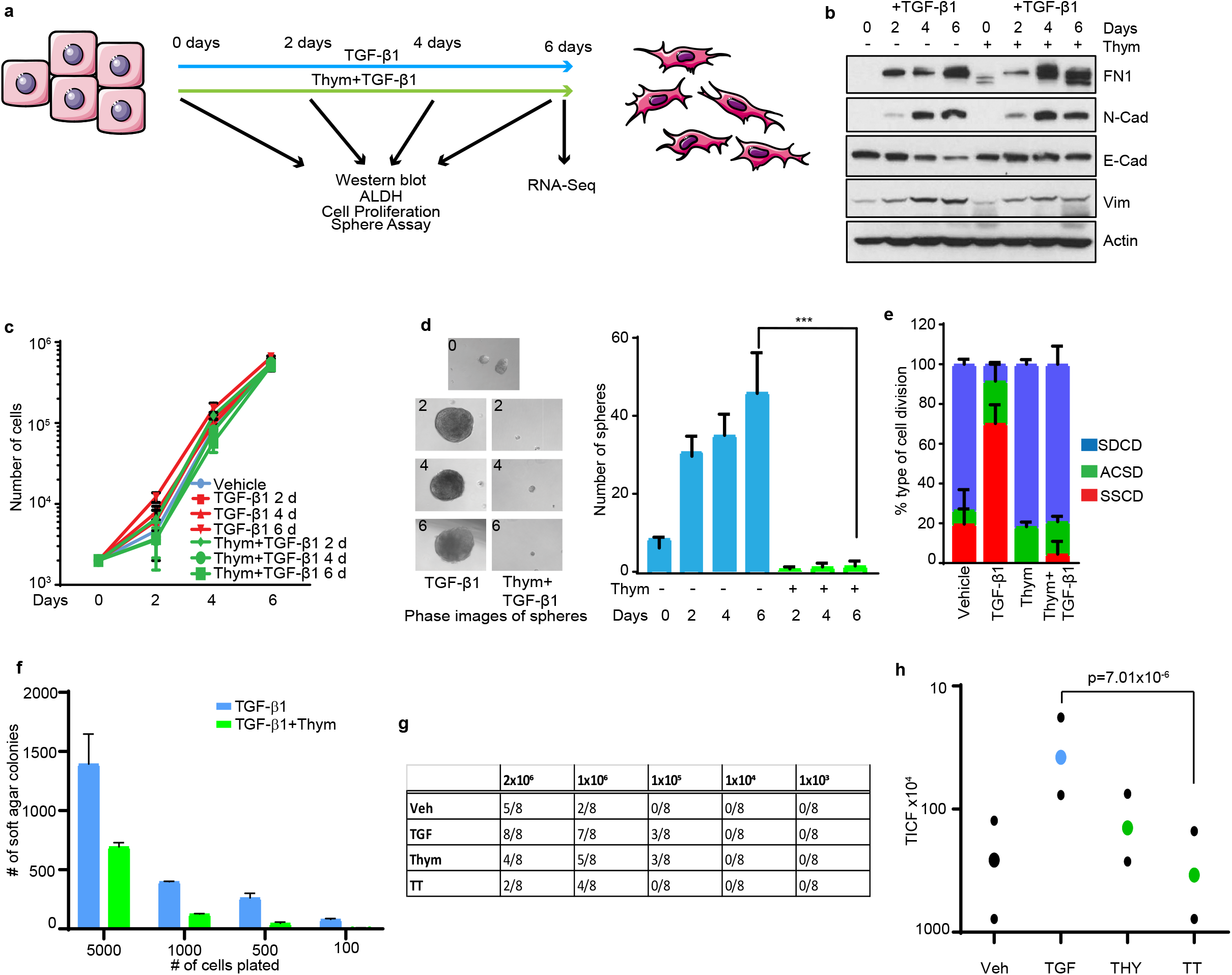
Cell division is required for EMT-induced stemness. **a**, Cartoon depicting the experimental timeline of TGF-β1 and thymidine treatments and indicated assays. **b**, Western blot analysis of markers associated with EMT in MCF10A cells treated with TGF-β1 alone or with thymidine at indicated times. **c**, Proliferation of MCF10A cells after 2, 4, and 6 days TGF-β1 treatment with or without thymidine (n=3). **d**, Number of spheres formed from 500 MCF10A cells following TGF-β1 treatment alone or with thymidine (n=4). **e**, Quantitation of SDCD, ACSD, and SSCD cell division after 6 days of TGF-β1 treatment with or without thymidine (n=3). **f**, Quantitation of limiting dilution soft-agar colony-forming ability after 6 days of TGF-β1 treatment with or without thymidine (n=3). **g**, Number of cells injected in each mouse and the number of mice per group that developed tumors. Prior to injection, MCF10A Ras cells were treated for 6 days with vehicle (Veh), TGF-β1 (TGF), thymidine (Thy), or TGF-β1 plus thymidine (TT). **h**, Number of tumor-initiating cells per treatment based on the data shown in panel f. The p values were calculated using chi-squared test.

To determine whether cell division inhibition affected in tumor initiating stemness we used MCF10A cells transformed with oncogenic H-ras (MCF10A-Ras), which renders the cells capable of forming tumors *in vivo*. We first tested whether the reduction in stemness when cell division is blocked during EMT is able to affect soft-agar colony formation. Accordingly, we treated MCF10A-Ras cells with both TGF-β1 and thymidine. Consistent with our other stem cell assays, these cells showed a reduced capacity to form colonies as compared to control cells treated with TGF-β1 alone (Fig. 3f). Next, we tested whether the reduction in colony formation in soft-agar correlated with the tumor initiating activity determined using a limiting dilution tumor initiation assay (LDTI). For this, we pretreated MCF10A-Ras cells with TGF-β1 and/or thymidine and injected 2×10^6^, 10^6^, 10^5^, 10^4^ and 10^3^ cells into the fourth mammary fat pad of NOD/SCID mice. Similar to the *in vitro* findings, the tumor-initiating cell frequency is significantly reduced from 1/430,000 in TGF-β1-treated cells to 1/3,430,000 (a 9.6-fold reduction) after blocking cell division with thymidine (Fig. 3 g and h). The significant reduction or absence of ALDH activity, lack of sphere formation, failure to switch cell division to SSCD, and reduced tumor-initiating cell frequency *in vivo*, in the absence of cell division, provide support for the conclusion that cell division is critical for gaining stemness during EMT.

## Cell division is not necessary to maintain the stem-cell properties

While these studies indicate that epithelial cells need to divide to acquire stem cell properties while undergoing EMT, it remains unclear whether continued cell division is necessary to maintain stem cell properties. To test this, we induced EMT in MCF10A cells by exposing them to TGF-β1 for 6 days and then exposed them to TGF-β1 and thymidine for an additional 6 days (Fig. 4a). Inhibition of cell division after the cells have undergone EMT neither affected the expression of EMT markers (Fig. 4b) nor proliferation (Fig. 4c) or stem-cell properties (Fig. 4d and Supplementary Fig. 4a). Similarly, inhibition of cell division did not affect the stem cell and mesenchymal properties in HMLER cells stably expressing Snail (HMLER-Snail) (Supplementary Fig. 4b-f). Furthermore, the tumor-initiating cell frequency in MCF10A-Ras cells induced to undergo EMT in response to TGF-β1 and then subjected to thymidine exposure was 1/1,480,000 as compared to 1/1,570,000 of the same cells without thymidine exposure. This difference is not statistically significant (Fig. 4e and f). These data strongly suggest that cell division is critical for acquiring stem cell properties during EMT but not for the maintenance of stemness in cells that have already undergone EMT.

**Figure 4.**
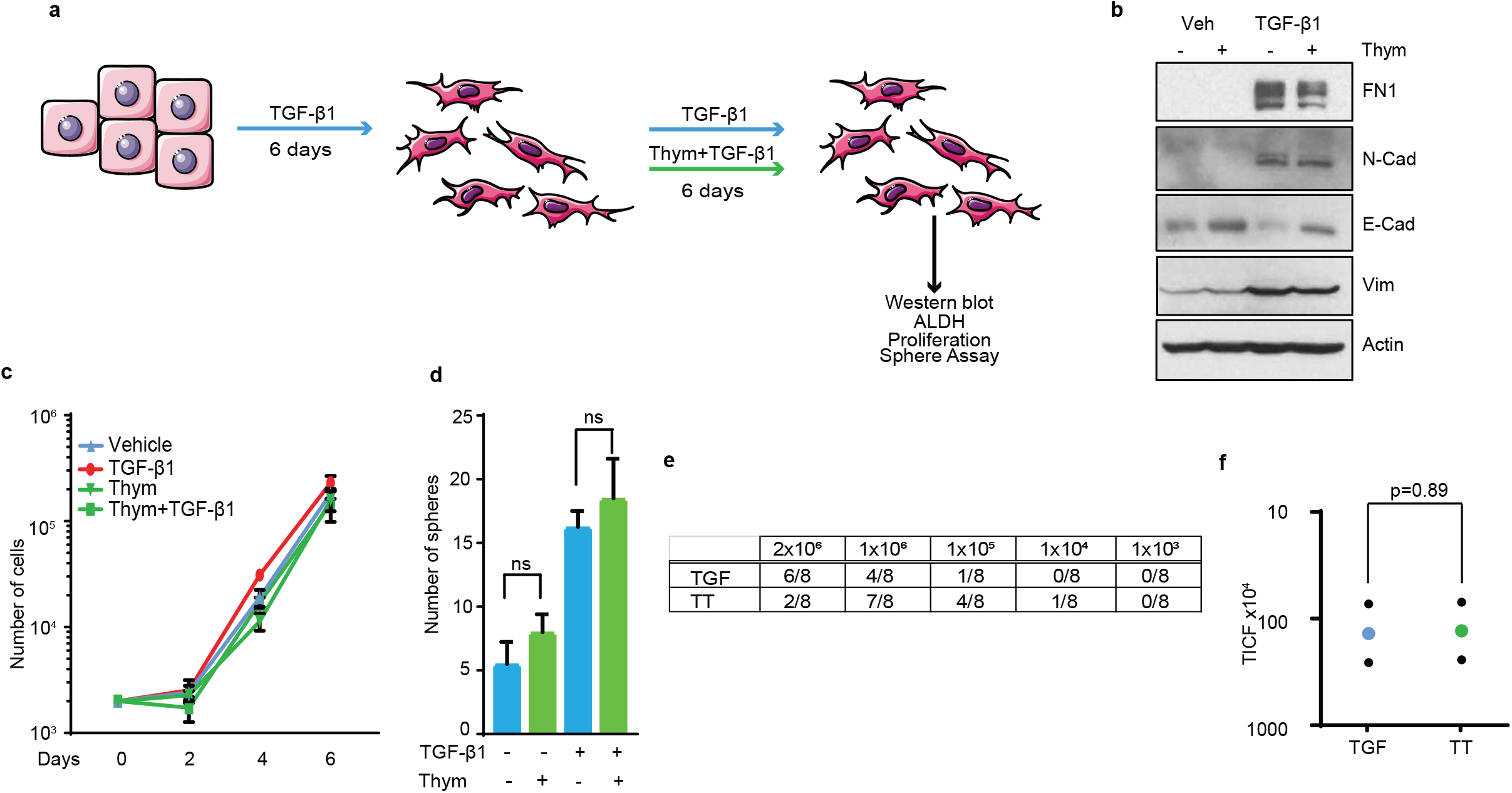
Cell division is not critical to maintain stemness of EMT cells. **a**, Cartoon of the experimental timeline of TGF-β1 and thymidine treatments and the assays performed. **b**, Western blot analysis of markers associated with EMT in MCF10A cells after 6 days of TGF-β1 exposure and 6 days TGF-β1 and thymidine exposure. **c**, Proliferation of MCF10A cells after TGF-β1 and/or thymidine exposure (n=3). **d**, Number of spheres formed by 500 MCF10A cells collected after indicated treatments (n=4). Error bars show standard deviation. **m**, Number of cells injected in each mouse and the number of mice per group that developed tumors. MCF10A Ras cells were pretreated for 6 days with TGF-β1 and 6 days with TGF-β1 (TGF) or with TGF-β1 and thymidine (TT). **n**, Number of tumor-initiating cells per treatment based on the data shown in panel m. The p values were calculated using chi-squared test.

## EMT-induced stem cell signature and not the mesenchymal signature predicts poor clinical outcome

Next, to identify the genes unique to EMT-induced stem cell properties, we performed RNA-seq analysis on 6-day TGF-β1-treated MCF10A cells with and without thymidine exposure (Supplementary Fig. 5a). Using multidimensional scaling of the RNA-seq data, we identified differentially regulated genes in response to TGF-β1 (Fig. 5a) and TGF-β1 plus thymidine treatment (Fig. 5b) relative to control cells treated with vehicle. Using gene set-enrichment analysis (GSEA), we found that genes associated with TGF-β1 signaling are upregulated in cells treated with TGF-β1 alone or in combination with thymidine, as expected (p = 0.001, Supplementary Fig. 5b,c). Analysis of genes and pathways selectively affected in response to cell division inhibition during TGF-β1-induced EMT identified a subset of genes enriched for stem cell-related genes and pathways (Fig. 5c-d). In contrast, genes that are unaffected by cell division inhibition are primarily enriched for mesenchymal-related genes (Fig. 5a and c; Supplementary Fig. 5d, e). Inhibition of cell division results in a significant reduction in stem cell (ES1) signature (NES=-1.31, p=0.007), but not mesenchymal genes (NES=2.09, NES=1.88 respectively) (Fig. 5c and d, Supplementary Fig. 5b-c). These data reiterate our experimental observation that stem cell programs dependent on cell division and not the mesenchymal programs during EMT.

**Figure 5.**
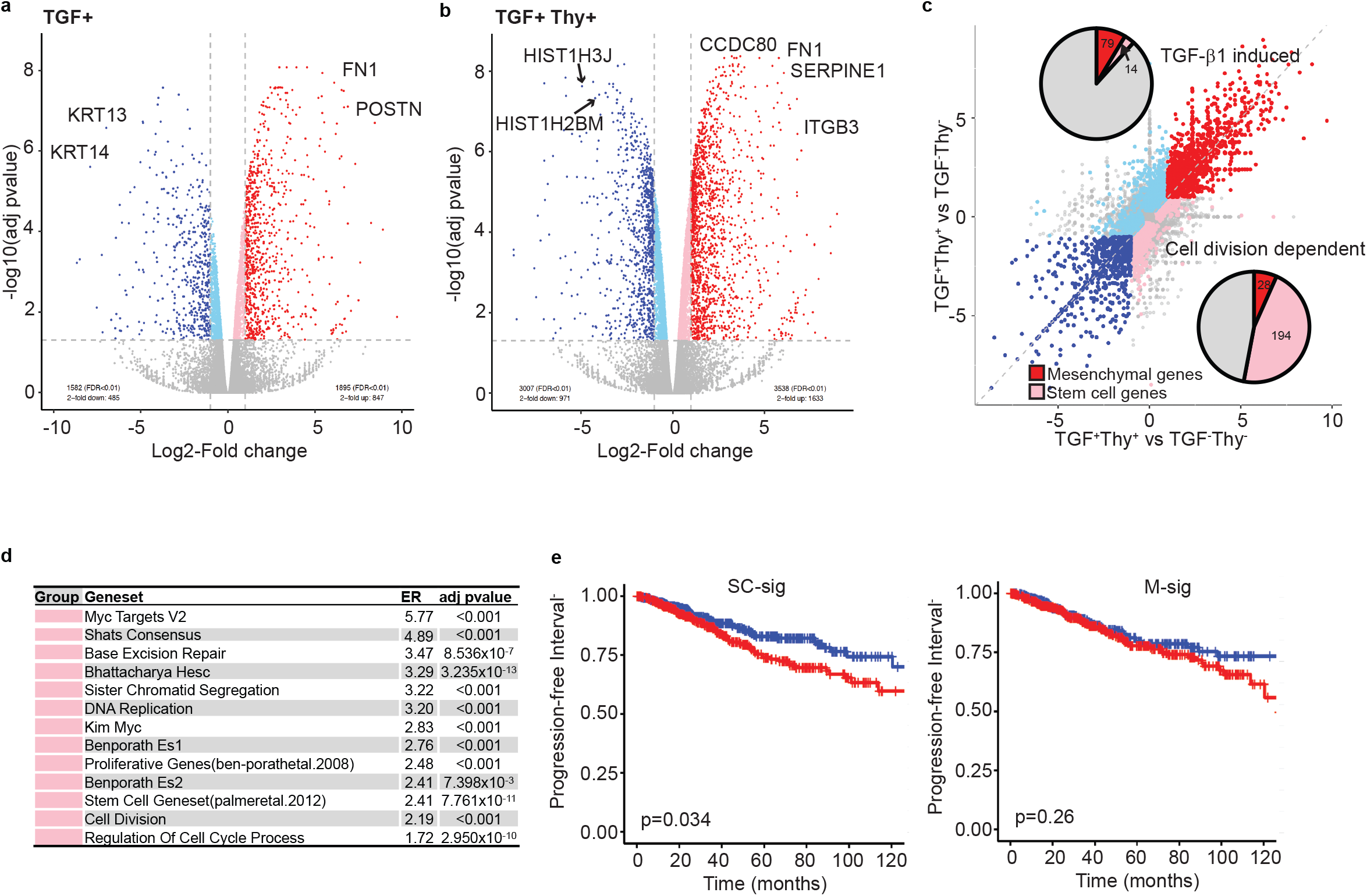
Inhibition of cell division affects the expression of genes associated with stemness. **a**, Differences in expression as Log2-fold change with TGF-β1 treatment compared to vehicle treatment. **b**, Differences in expression as Log2-fold change with TGF-β1 plus thymidine treatment compared to vehicle treatment. **c**, Comparison of gene expression difference between TGF-β1 alone and TGF-β1 plus thymidine treatment. Pie charts reflect the representation of mesenchymal-related (red) and stem cell-related genes in the TGF-β1-induced and cell division-dependent (light red) gene groups. **d**, Pathway enrichment analysis within the genes selectively affected by TGF-β1 treatment (red) and the genes selectively affected by cell division inhibition during TGF-β1-induced EMT (light red). **e**, Kaplan-Meier plot of PFI for breast cancer patients with high and low levels of SC-sig, and M-sig genes.

We then used the mesenchymal-associated genes upregulated within the TGF-β1-induced group (red) to develop a mesenchymal gene signature (M-sig). Likewise, using the stem cell-associated genes upregulated within the cell division-dependent group, we established a stem cell signature (SC-sig). Since EMT is essential in cancer progression, recurrence, and resistance to therapy ^4,16,17,18,19^, we determined the clinical association of M-sig and SC-sig with survival using the large TCGA breast cancer patient data set encompassing all types of breast cancer. Remarkably, our SC-sig effectively predicted progression-free interval, disease-specific survival, and overall survival of these breast cancer patients, while the EMT-induced M-sig could not (Fig. 5e, Supplementary Fig. 6a-d). These data indicate that the EMT-induced stem cell program, but not the mesenchymal program, are the drivers of disease progression during EMT.

## Discussion

Our studies show that EMT-inducing signals can trigger the mesenchymal program in cancer cells without a need for cell division; still, they need to undergo cell division to gain stem cell properties during EMT. TGF-β1 is able to induce the mesenchymal program without the need for cell division suggesting that the mesenchymal program may require only signal transduction signaling without any cellular reprogramming. The requirement of cell division for the TGF-β1 or Snail to induce stem cell properties suggests that cancer cells need to modify the genome gradually to take on the stem cell phenotype, probably via epigenetically altering the genome.

The fact that the cells that have undergone an EMT retain the stem cell properties without relying on further cell division reiterates that once the genome is epigenetically reprogrammed to become a stem-like cell, it keeps those properties independent of cell division. These findings also suggest that blocking cell division can inhibit the acquisition of stem cell properties; simultaneously, inhibitors of the epigenetic program may revert the stem cell properties in cells that have undergone EMT and acquired stem cell properties. Clinically, it has been challenging to distinctly identify the aggressive tumors rich in cancer cells displaying mesenchymal and stem cell properties gained via EMT from the other tumors that are less aggressive and enriched with mostly fibroblasts. Our new EMT-induced stem cell signature SC-sig can help identify EMT-induced, aggressive, metastasis-prone tumors rich in stem cells and may help separate the above two groups of tumors.

## Methods

### Cell lines, cell culture, reagents, and treatments

MCF10A cells (ATCC) were cultured in DMEM/F12 containing 5% horse serum, 10 µg/ml insulin (Sigma), 1 mg/ml hydrocortisone (Sigma), 20 ng/ml EGF, 100 ng/ml cholera toxin (Sigma), 5% penicillin/streptomycin (Gibco/Life Technologies). HMLER Snail-ER and HLMER-Snail (a gift from Robert Weinberg) cells were grown in 50% MEGM (Lonza) and 50% DMEM/F12 50:50 (Corning) supplemented with bovine pituitary extract (Lonza), 10 µg/ml insulin (Sigma), 1 mg/ml hydrocortisone (Sigma), 5% penicillin/streptomycin (Gibco/Life Technologies). TGF-β1 (Novus Biologicals) was used at 5 ng/ml final concentration, thymidine (Sigma) at 500 µg/ml, lovastatin (Sigma) at 20 µM, mevalonolactone (Sigma) at 100 mg/ml, and 4-Hydroxytamoxifen (Sigma) was used at a final concentration of 100 nM.

### Western blot analysis

Proteins were extracted using RIPA buffer (Sigma) containing cOmplete™ protease inhibitor cocktail (Roche) and PhosphoStop phosphatase inhibitors (Roche). Protein concentrations were determined using BIORad Bradford assay. Aliquots of 30 µg of protein were separated SDS/PAGE, transferred to nitrocellulose membranes, probed sequentially with antibodies of interest, *viz*., Actin (Sigma), Fibronectin (BD Biosciences), Vimentin (Thermo Fisher), N-Cadherin (BD Biosciences), E-Cadherin (BD Biosciences), and HRP-labelled secondary antibodies with chemiluminescence was used for detection.

### Mammosphere assay

Cells were treated with TGF-β1 (or tamoxifen) and/or thymidine for the indicated number of days. Cells were then trypsinized, counted on a hemocytometer, and 500 cells were plated per well in an ultra-low attachment 96-well plate in 100 µl of the mammosphere media (MEGM media with 1% methylcellulose containing 10 ng/ml EGF, 20 ng/ml FGF, and 4 µg/ml heparin). Fresh mammosphere media was replaced every 3 days, and spheres were counted after 14 days. Experiments were performed in quadruplicates.

### ALDH assay

Cells were treated with TGF-β1 and/or thymidine for the indicated number of days. Cells were then trypsinized, and 100,000 cells were tested in the ALDefluor assay as per the manufacturer’s protocol (Stem Cell Technologies). Cells were incubated with ALDH reagent for 45 min and were analyzed using a BD Accuri. Experiments were performed in triplicate.

### Cell-cycle phase analysis

MCF10A cells were treated with TGF-β1 and/or thymidine for the indicated times and fixed in 70% ethanol overnight. Cells were treated with RNase, stained with propidium iodide, and DNA content was analyzed using the LSR Analyzer. Experiments were performed in triplicate.

### Cell division analysis

Cells were treated with TGF-β1 for the indicated amount of time and then treated with 100 nM of nocodazole for 14-16 h to arrest cells in G2. Mitotic cells were collected and incubated in the growth medium for 90 min to induce cell division. Cells were then fixed in 4% paraformaldehyde for 15 min and spotted on slides using a cytospin centrifuge. The slides were blocked with 5% normal goat serum containing 0.1% Triton X-100.

Immunofluorescent staining was carried out by incubating the samples in anti-NUMB antibody (Cell Signaling) overnight at 4 °C and detected with Alexa-488-labeled secondary antibody (Life Technologies). Images were taken using a Zeiss inverted fluorescent microscope, and types of cell division events were counted and charted in a graph. Experiments were performed in triplicate.

### Anchorage-independent growth assays

Anchorage-independent growth assays were performed after 6 days of treatment with vehicle, TGF-β1, thymidine or TGF-β1 and thymidine by plating 5000, 1000, 500 or 100 cells in 0.35% SeaPlaque GTG Agar (LONZA) in MCF10A medium on top of a 0.7% agar base in MCF10A medium. Colonies were scanned using the Cytation 3 and counted after 2 weeks. All experiments were performed in triplicate, and results reported as average colony number ± SD.

### Limiting dilution tumor initiation assays

MCF10A-Ras cells were treated for 6 days with vehicle, TGF-β1, thymidine or TGF-β1 and thymidine before injecting into the 4^th^ mammaray gland of NOD/SCID mice. Celles were mixed 50/50 with Matrigel and either 2×10^6^, 1×10^6^, 1×10^5^, 10x^4^ or 1×10^3^ cells were injected bilaterally in 4 mice per dilution group (8 data points). After 6 weeks tumors were measured and tumors bigger than 5mm were included in the anlysis to calculate tumor-initating cell frequency.

### RNA extraction and RNA-seq analysis

MCF10A cells were treated with vehicle (control), TGF-β1, or thymidine or TGF-β1 plus thymidine for 6 days. Cells were lysed on the plate, and RNA was isolated using a Qiagen RNeasy kit. Two samples were analyzed per condition. RNA concentrations were determined using a Nanodrop spectrophotometer. RNA quality was assessed using Agilent Bioanalyzer. RNA-seq libraries were prepared from 100 ng of total RNA using RNA HyerPrep kit with RiboErase (Kapa Biosystems). Sequencing was performed at the MD Anderson core facility equipped with an Illumina HiSeq 4000 instrument. Sequencing quality was evaluated using the FastQC software (http://www.bioinformatics.babraham.ac.uk/projects/fastqc/).

mRNA sequencing reads were mapped to the human genome hg19 by STAR (version 2.5.3) using the 1-pass model. Hg19 sequence and RefSeq annotation were downloaded from the UCSC table browser (03/29/2017). RSEM (version 1.2.31) was used to quantitate transcript levels. The estimated counts were normalized by upper-quartile normalization implemented in edgeR (v3.7) and log2-transformed. Normalized data were used for downstream analysis.

### Multidimensional scaling plot

Multidimensional scaling analysis was performed on log2-transformed normalized mRNA counts using the MDS function implemented in the edgeR package. The log2-fold-change distances obtained from multidimensional scaling between the samples in dimension 1 (dim1) and dimension 2 (dim2) were plotted. Data were fit using the LOESS (locally weighted smoothing) function. Principal component analysis was performed using factoextra (v1.0.6) implemented in R. The first principal component (PC1) explained 33.9% of the variability, whereas the second principal component (PC2) explained 24.5% of the variability in gene expression between MCF10A samples.

### Differentially expressed genes (DEG) analysis

Genes differentially expressed in TGF-β1-treated and untreated MCF10A cells were identified using quasi-likelihood linear modeling implemented in edgeR (v3.7). Using the same method, DEG between TGF-β1 and thymidine-treated cells and cells treated only with TGF-β1 were identified. The false discovery rate was calculated using the Benjamini and Hochberg method. Statistical analysis was performed by moderated t-test implemented in the limma package in edgeR. Genes with adjusted p-value ≤ 0.05 and fold change ≤ −2 and ≥ 2 were considered as significant. The heatmap was prepared using ComplexHeatmap (v2.1.0) library in R.

### Gene-set enrichment analysis (GSEA)

Before enrichment analysis, the replicates were averaged at the gene level. Enrichment analysis was performed using WebGestaltR [13] using gene ontology, hallmarks gene sets downloaded from MSigDB (v6.2), and the stem cell gene set [14]. The gene sets were considered significant at an adjusted p-value ≤ 0.05. Enrichment ratios are given for each gene set. GSEA was performed on differentially expressed genes using fgsea (v1.6.0). Single-sample GSEA (ssGSEA) was performed using the GSVA package (v1.28.0) using the hallmarks gene set. The normalized enrichment score for gene sets was scaled and compared between samples.

### Overlap with known EMT and stem-cell gene sets

We downloaded the hallmark of the EMT gene set (200 genes) and the TGF-β1 signaling gene set (54 genes) from MSigDB (v6.2). For stem cell-related genes, we downloaded Shats Consensus and Benporath ES1 (overexpressed in human embryonic stem cells) gene sets corresponding to 80 and 361 genes, respectively [14]. The fibroblast-associated gene set was retrieved from gene ontology terms *GO regulation of fibroblast proliferation* and *GO regulation of fibroblast migration* consisting of 81 and 28 genes, respectively.

### Patient data

We downloaded the cancer genome atlas (TCGA) RNA-seq normalized count data and clinical characteristics for breast cancer (http://gdac.broadinstitute.org/, January 2016 version). Clinical data with overall survival time, vital status, and tumor stage were considered in the analysis. Curated survival endpoints were retrieved from XENA (https://xenabrowser.net/) [15]. The clinical endpoints available are overall survival (OS), disease-specific survival (DSS), disease-free interval (DFI), and progression-free interval (PFI).

### Signature score

In this study, we identified mesenchymal-enriched (79), stem-cell enriched (194), and both mesenchymal and stem cell-enriched (273) differential gene signatures. For comparison, the EMT gene set from MSigDB with 200 genes was used. The ssGSEA was performed on TCGA patient RNA-seq data (normalized and log2-transformed) using these gene signatures.

### Survival analysis

Survival for each patient was calculated as the time from date of diagnosis to death or last contact, progression-free interval time, disease-free interval time, and disease-specific time. A normalized enriched score (NES) of each signature gene set derived was categorized into low and high expression based on the mean. Univariate Cox proportional hazards models were fitted to overall survival, disease-specific survival, disease-free interval, and progression-free interval endpoints for patients to calculate the hazard ratios using coxph function in the Survival package (version 2.44). The p-values were determined using a log-rank test. The hazard ratio [*exp(cox coefficient)*] was used to compare survival in signature M-sig, SC-sig, EMT-sig and Hallmark EMT-MSigDB geneset. Boxplots used for hazard ratio visualization across cancer cohorts were prepared using R (version 3.5.2). Cox coefficients (ß) for gene sets across cancer cohorts are shown in dumbbell plots with ß>2 considered as 2 and ß< -2 considered as - 2 for visualization purposes.

### Statistical analyses

The statistical analyses used are specified in the figure legends. The t-tests and survival analyses on patient data were performed using t.test and survival functions in R (Bioconductor, version 3.5.2). The experimental dataset was compared with t-tests using GraphPad Prism. The p values less than 0.05 were considered to be statistically significant.

## Acknowledgments

This work was supported by a grant from the NIH/NCI to SAM (2R01CA155243) and translational genomics and precision medicine in cancer training program T32 award (5T32CA217789-04) to MC. In addition, we thank Dr. Jeffrey Rosen, Baylor College of Medicine, for his valuable input and support.

## Author contributions

P.d.H. performed experiments, data collection, data processing, experimental and conceptual design, and manuscript preparation. S.V. performed RNA-seq data analysis and gene signature development, patient survival data processing and analysis, and manuscript preparation. M.C. performed experiments and data processing. R.J. performed experiments and manuscript preparation. A.D performed analysis and data processing. T.Z. and M.P. performed experiments. C.F. performed RNA-seq library preparation and quality assurance. W.F.S. supervised RNA-seq library preparation and quality assurance. R.S. performed experimental and conceptual design, project management, generation and maintenance of all protocols required for compliance for this project, and manuscript preparation. S.A.M. conceived the idea and oversaw experimental and conceptual designs, experiments and data processing, and manuscript preparation.

## Competing Interests Statement

The authors declare no competing interests.

## Notes

### Competing Interest Statement

The authors have declared no competing interest.

