## Supplemental figures for "Acquisition of cancer stem cell properties during EMT requires cell division"

Supplemental Figure 1

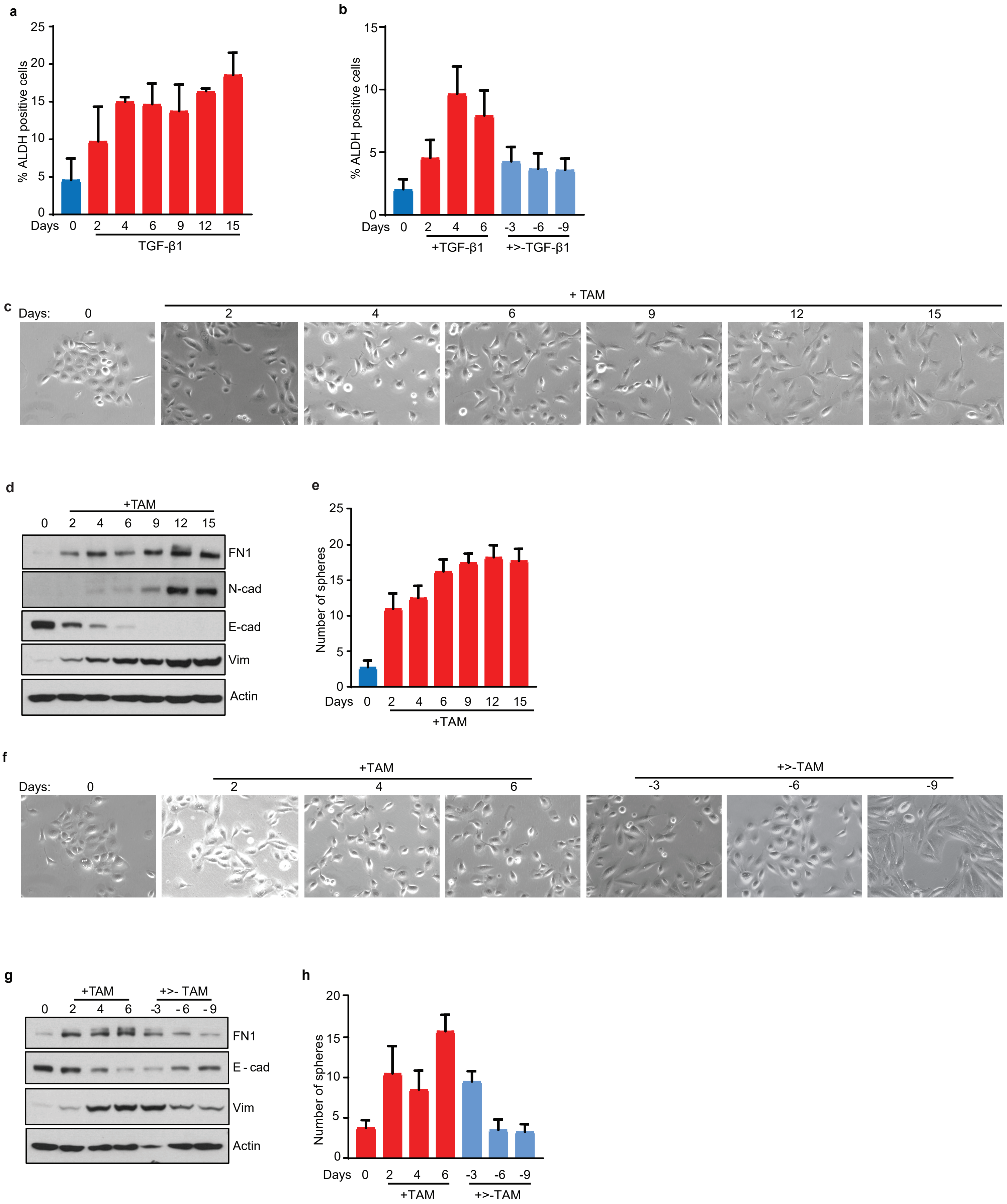

Supplemental Figure 2

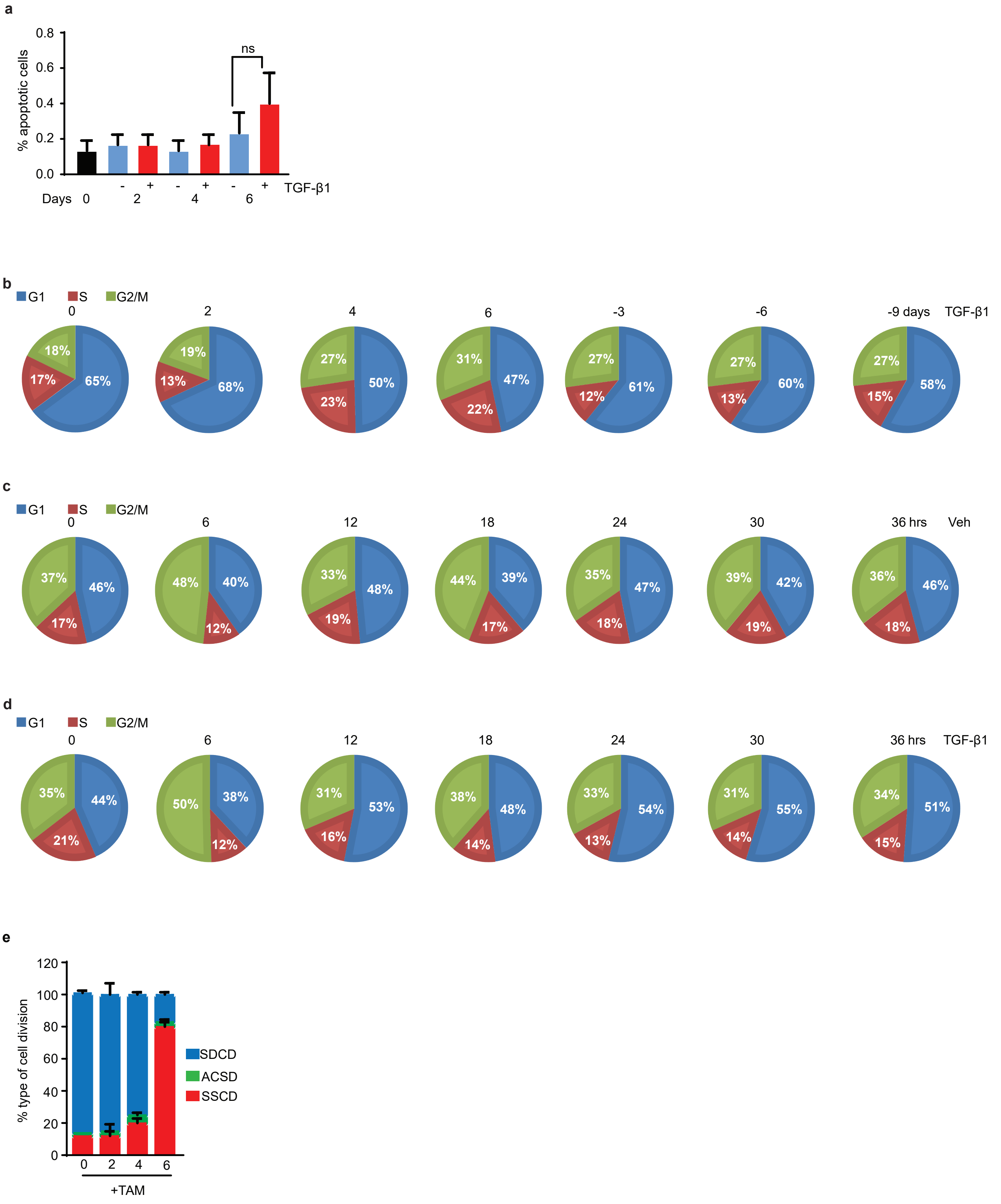

Supplemental Figure 3

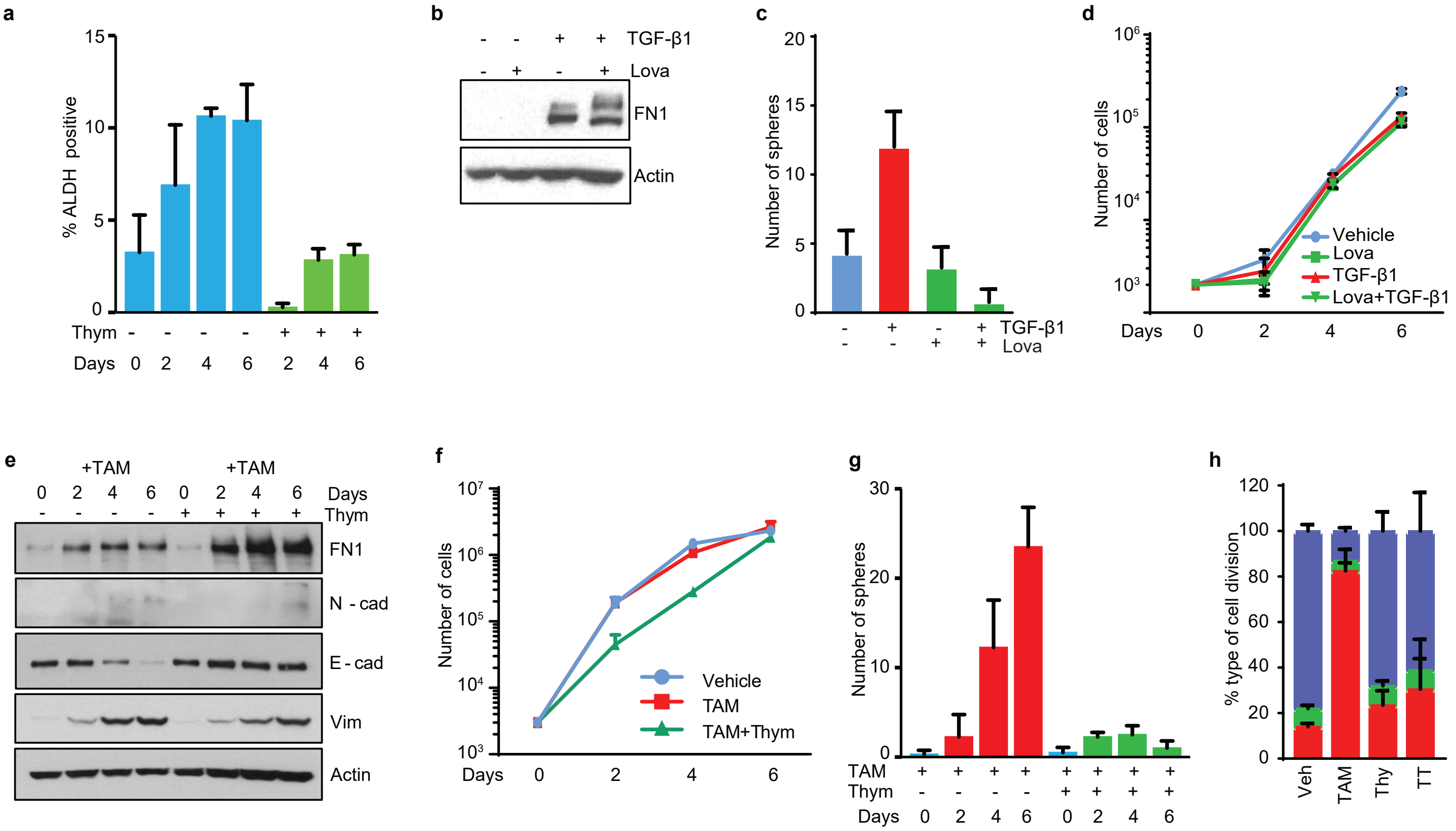

Supplemental Figure 4

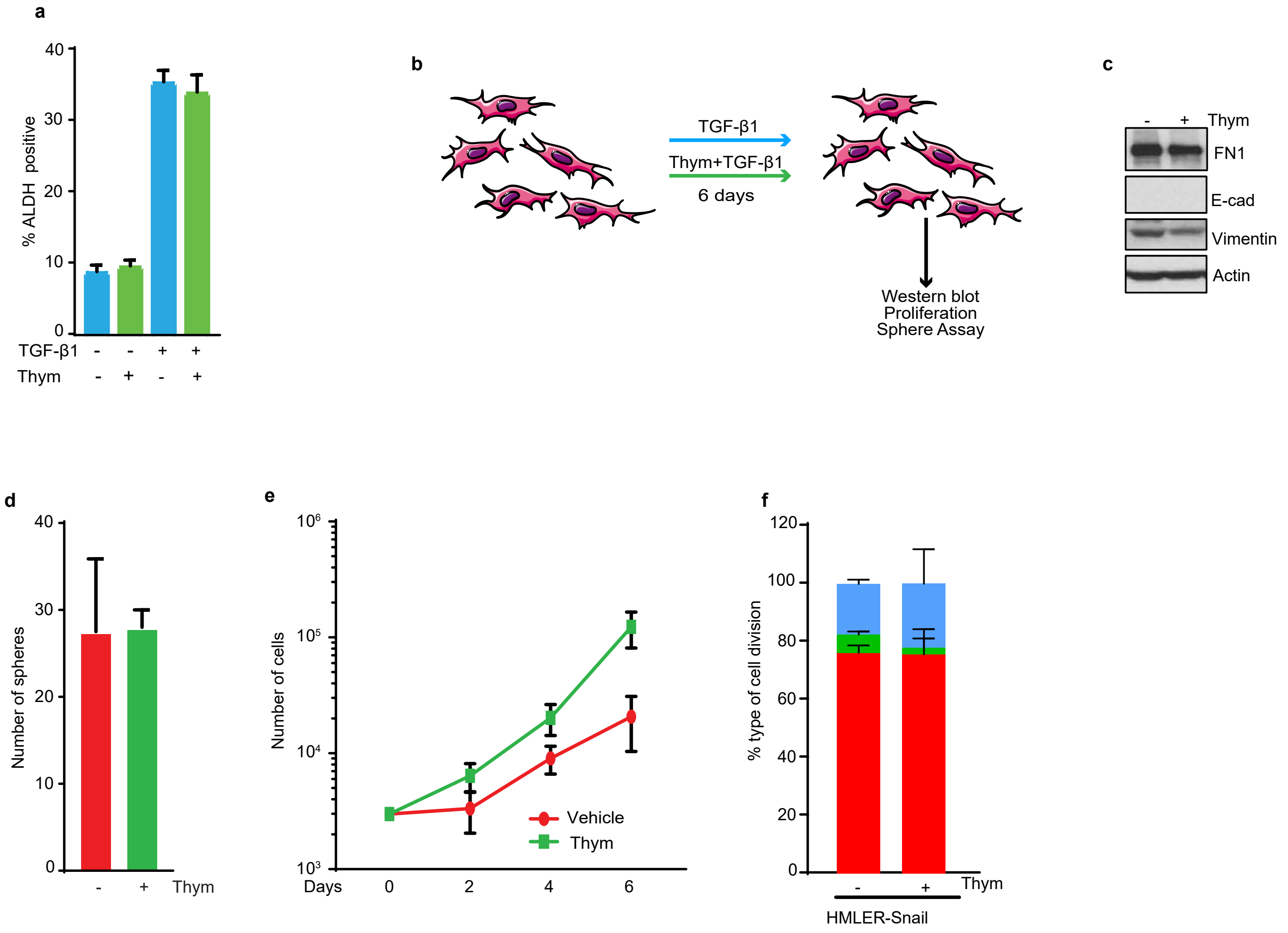

### Supplemental Figure 5

a

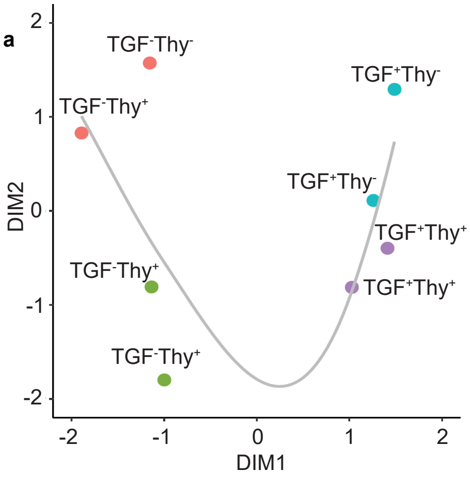


b

| Gene Ontology | TGFB+ NES | TGFB+ pvalue | TGFB+Thy+ NES | TGFB+Thy+ pvalue |
|---|---|---|---|---|
| Epithelial Mesenchymal Transition | 2.095 | 1.51x10$^{-3}$ | 1.893 | 1.26x10$^{-3}$ |
| TGF Beta Signaling | 1.684 | 1.74x10$^{-3}$ | 1.755 | 1.50x10$^{-3}$ |
| Stem Cell Markers (TFs, CSCdb) | 1.293 | 3.13x10$^{-2}$ | 1.013 | 0.416 |
| Benporath ES1 | 0.785 | > 0.9 | -1.321 | 7.04x10$^{-3}$ |
| Cell Division | 0.781 | > 0.9 | -1.207 | 2.73x10$^{-2}$ |
| Nuclear Chromosome Segregation | 0.683 | > 0.9 | -1.421 | 9.52x10$^{-3}$ |
| Proliferative Genes(ben-porathetal.2008) | 0.663 | > 0.9 | -1.232 | 2.38x10$^{-2}$ |
| Mitotic Cell Cycle | -0.682 | > 0.9 | -1.316 | 1.75x10$^{-2}$ |
| Mitotic Nuclear Division | -0.757 | > 0.9 | -1.503 | 6.45x10$^{-3}$ |
| Cellular Lipid Metabolic Process | -1.147 | 1.49x10$^{-2}$ | 0.829 | > 0.9 |

c

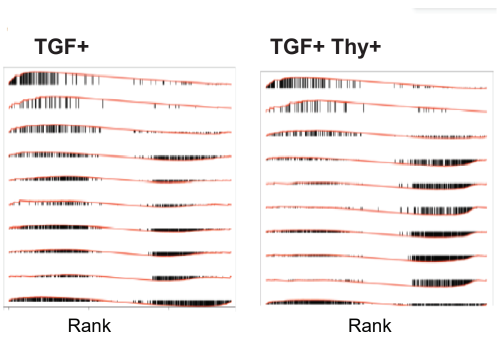


d

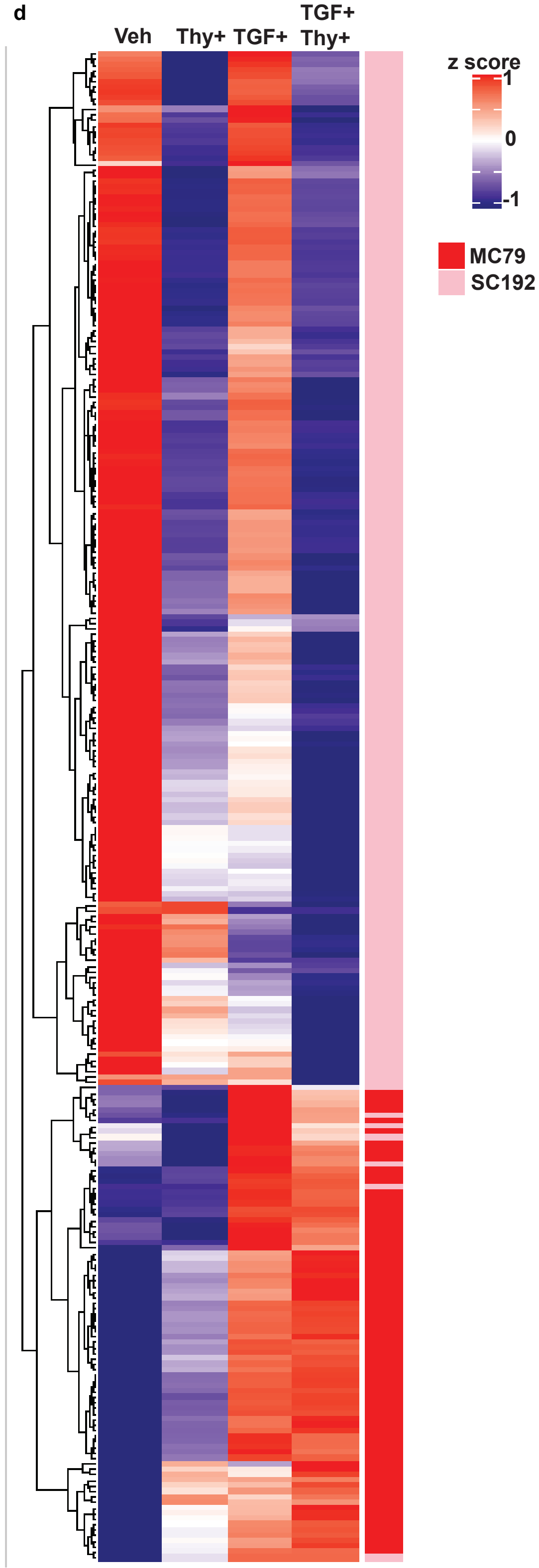


e

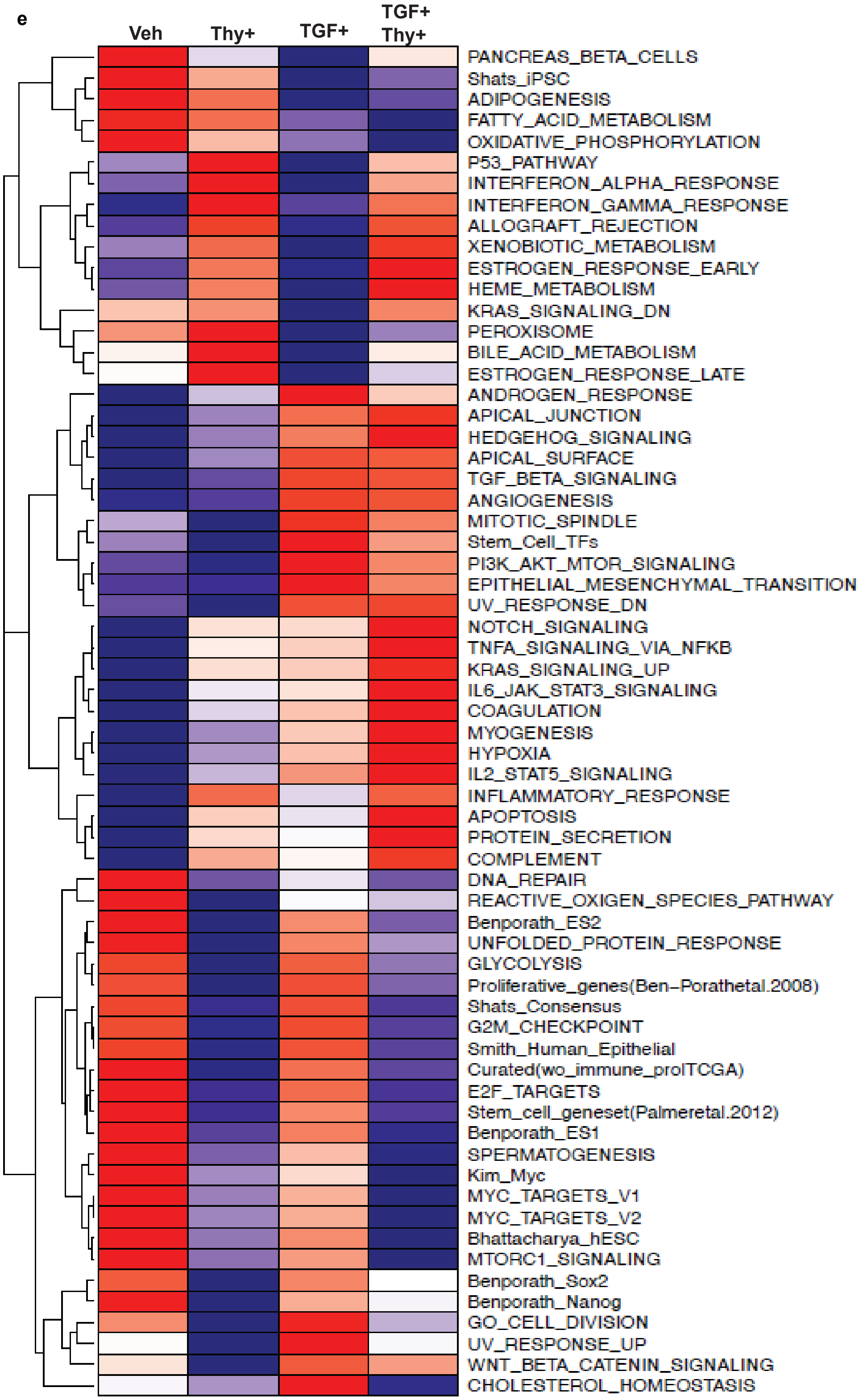

Supplemental Figure 6

a

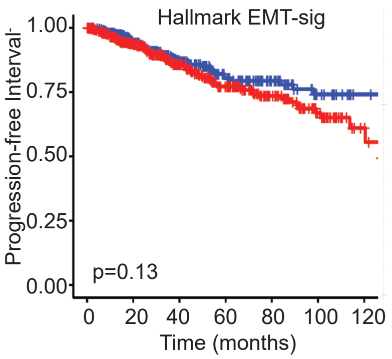


b

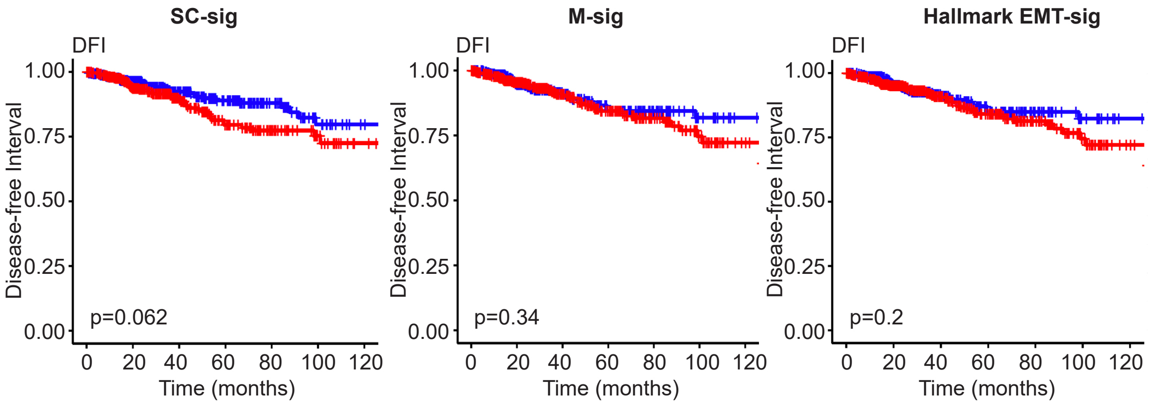


c

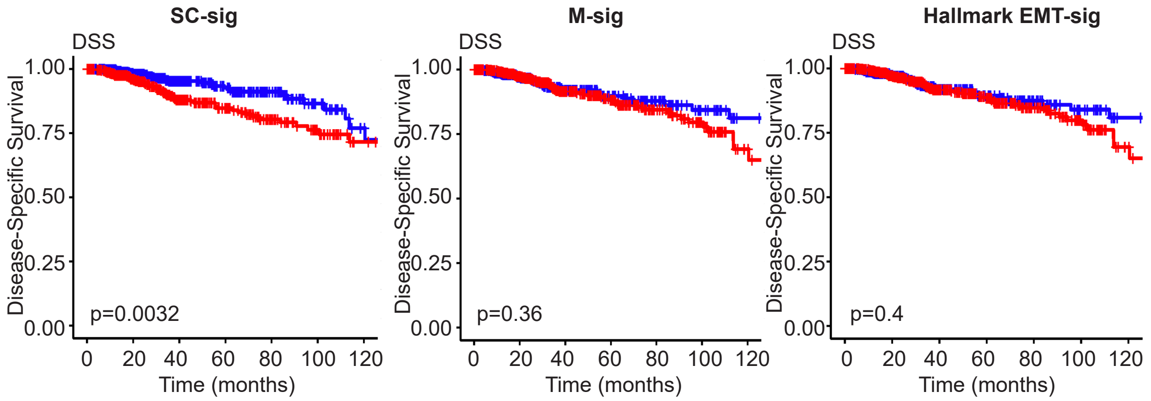


d

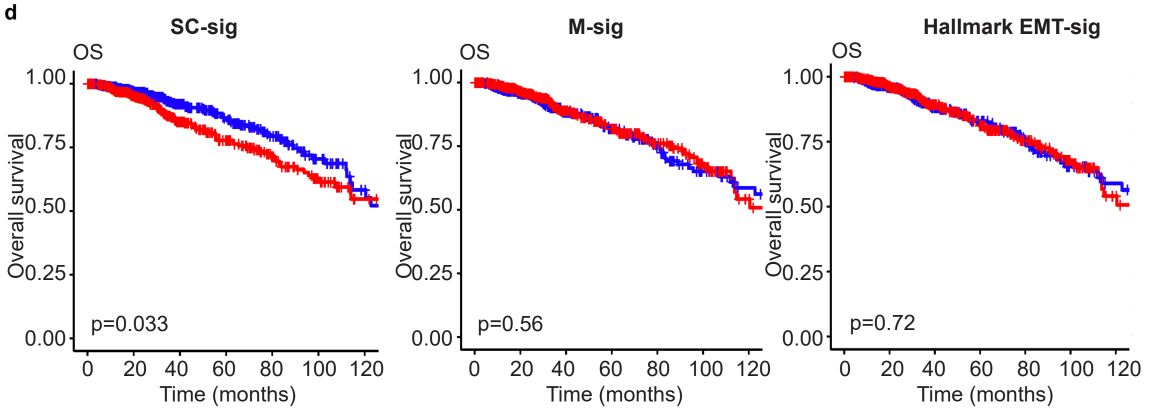

**Supplementary Figure 1 | Mesenchymal and stem cell properties induced in HMLER-Snail-ER cells are reversible.** **a,** Percentage of MCF10A cells with ALDH activity during fifteen days of TGF-β1 exposure (n=3). **b,** Percentage of MCF10A cells with ALDH activity during 6 days of TGF-β1 exposure and 9 days of withdrawal (n=3). **c.** Phase-contrast imaging of HMLER Snail-ER cells at indicated days during a 15-day tamoxifen exposure. **d,** Western blot analysis of markers associated with EMT at indicated days during tamoxifen exposure. **e,** Number of spheres formed from 500 HMLER Snail-ER cells collected on indicated days during tamoxifen treatment (n=4). **f,** Phase-contrast imaging of HMLER Snail-ER cells during 6-day tamoxifen exposure and 9-day withdrawal. **g,** Western blot analysis of markers associated with EMT during 6 days of tamoxifen exposure and 9 days of withdrawal. **h,** Number of spheres formed in samples of 500 HMLER Snail-ER cells collected on indicated days during tamoxifen treatment and withdrawal (n=4). Error bars reflect standard deviation.

**Supplementary Figure 2 | TGF-β1 does not significantly affect proliferation or apoptosis in MCF10A cells.** **a,** Apoptotic index of MCF10A cells exposed to TGF-β1 or vehicle for 2, 4, and 6 days, as measured by propidium iodide staining (n=3). **d,** Cell-cycle distribution of MCF10A cells after 2, 4, and 6 days of TGF-β1 or vehicle exposure (n=3). **c,** Cell-cycle progression of MCF10A cells treated with TGF-β1 or vehicle for the indicated number of hours (n=3). **d,** Quantitation of cell division type undergone by HMLER Snail-ER cells after tamoxifen exposure for 6 days (n=3). Error bars reflect standard deviation.

**Supplementary Figure 3 | Cell division inhibitors block the development of stemness but not mesenchymal transition.** **a.** Percent cells with ALDH activity during treatment with TGF-β1 alone or with thymidine for 6 days, as normalized to control (n=3). **b,** Western blot analysis of markers associated with EMT in MCF10A cells treated/untreated with TGF-β1 and with/without lovastatin (Lova). **c,** Number of spheres formed in samples of 500 MCF10A cells following 6 days of treatment with TGF-β1 or lovastatin or both (n=4). **d,** Proliferation of MCF10A cells after 6 days of treatment with TGF-β1 and/or lovastatin (n=3). **e,** Expression of markers associated with EMT in HMLER-Snail-ER cells during 6-day tamoxifen and/or thymidine exposure. **f,** Proliferation of HMLER-Snail-ER cells during 6-day treatment with TGF-β1 and/or thymidine (n=3). **g,** Number of spheres formed in samples of 500 HMLER-Snail-ER cells following tamoxifen and/or thymidine exposure (n=4). **h,** Quantitation of cell division type of HMLER-Snail-ER cells after tamoxifen and/or thymidine exposure for 6 days (n=3). Error bars reflect standard deviation.

**Supplementary Figure 4 | Stem-cell properties and type of cell division are not affected in EMT cells treated with thymidine**. **a,** Percent cells normalized to control with ADHD activity after TGF-β1 and thymidine exposure (n=3). **b,** Cartoon of the experimental design of thymidine-treated cells undergone EMT. **c**, Western blot analysis of markers associated with EMT in HMLER-Snail cells treated with or without thymidine. **d,** Number of spheres formed from samples of 500 HMLER-Snail cells treated with or without thymidine (n=4). **e,** Proliferation of HMLER-Snail cells with or without thymidine treatment (n=3). **f,** Quantitation cell division type of HMLER-Snail cells with or without thymidine treatment (n=3). Error bars reflect standard deviation.

**Supplementary Figure 5 | Changes in MCF10A cell gene expression upon treatment with TGF-β1 and/or thymidine.** **a**, Principal component analysis of RNA-seq data of vehicle-treated ($TGF^{-}Thy^{-}$), TGF-β1-treated ($TGF^{+}Thy^{-}$), thymine-treated ($TGF^{-}Thy^{+}$), and TGF-β1 plus thymine-treated ($TGF^{+}Thy^{+}$) MCF10A cells. **b,c,** Normalized enrichment scores for RNA-seq data from cells treated with TGF-β1 alone, thymidine alone, or TGF-β1 plus thymidine compared to that of vehicle-treated cells. **d,** Heatmap of expression (z score) of mesenchymal signature (M-sig) and stem cell signature (SC-sig) genes in cells treated with TGF-β1 and/or thymidine. **e,** Heatmap of z scores of the differentially expressed pathways in cells treated with TGF-β1 and/or thymidine.

**Supplementary Figure 6 | Stem cell signature genes are predictive of survival in breast cancer patients.** **a,** Kaplan-Meier plot of disease-free survival (PFI) for Hallmark EMT-sig in breast cancer. **b,** Kaplan-Meier plot of disease-free survival (DFI) for SC-sig, M-sig, and Hallmark EMT-sig in breast cancer. **c,** Kaplan-Meier plot of disease-specific survival (DSS) for SC-sig, M-sig, and Hallmark EMT-sig in breast cancer. **d,** Kaplan-Meier plot of overall survival (OS) for SC-sig, M-sig, and Hallmark EMT-sig in breast cancer.
